# People can learn new walking patterns without walking

**DOI:** 10.1101/2020.04.02.021949

**Authors:** Christine N. Song, Jan Stenum, Kristan A. Leech, Chloe Keller, Ryan T. Roemmich

## Abstract

Humans can learn many new walking patterns. People have learned to snowshoe up mountains, racewalk marathons, and march in precise synchrony. But what is required to learn a new walking pattern? Here, we demonstrate that people can learn new walking patterns without actually walking. Through a series of experiments, we observe that stepping with only one leg can facilitate learning of an entirely new walking pattern (i.e., split-belt treadmill walking). We find that the nervous system learns from the relative motion between the legs – whether or not both legs are moving – and can transfer this learning to novel gaits. We also show that locomotor learning requires active movement: observing another person adapt their gait did not result in significantly faster learning. These findings reveal that people can learn new walking patterns without bilateral gait training, as stepping with one leg can facilitate adaptive learning that transfers to novel gait patterns.

## INTRODUCTION

People are capable of learning and storing many different walking patterns. The capacity for locomotor learning allows us to develop repertoires of walking patterns suitable for nearly any environment that we encounter. We tip-toe in a quiet room, trudge through snow, and limp in response to pain or injury. Locomotor learning also forms the basis of many rehabilitation approaches that aim to improve gait patterns following neurologic or musculoskeletal impairment (Reisman et al., 2010; Roemmich and Bastian, 2018).

Locomotor learning is not a singular process. People use a variety of mechanisms – e.g., reinforcement learning (Hasson et al., 2015), adaptation (Reisman et al., 2005), and strategic learning (French et al., 2018) – to learn new walking patterns. These learning mechanisms have specific underlying neural substrates (Glimcher, 2011; Morton and Bastian, 2006; Roemmich and Bastian, 2018; Taylor and Ivry, 2014) and can be leveraged in isolation or in combination (Cherry-Allen et al., 2018; Statton et al., 2016). The diversity of available learning mechanisms allows the nervous system to use information from different sources (e.g., feedback, error, instruction) to change walking patterns.

Adaptation in particular has emerged as an increasingly popular approach for facilitating locomotor learning (Reisman et al., 2010; Torres-Oviedo et al., 2011). This is because adaptation occurs relatively quickly (i.e., within minutes), is not dependent on external feedback or instruction, and causes gait changes that persist immediately after training (i.e., “aftereffects”) (Dietz et al., 1994; Jensen et al., 1998; Prokop et al., 1995; Reisman et al., 2005). Adaptation is often studied using a split-belt treadmill paradigm where each leg walks at a different speed (Dietz et al., 1994; Reisman et al., 2005). Over time, people adapt to walk with symmetric steps despite the asymmetry in belt speeds (Dietz et al., 1994; Reisman et al., 2005; Zijlstra and Dietz, 1995), and learning is quantified by measuring the gradual decrease in step length asymmetry. Prior studies have used this paradigm to understand how various clinical conditions affect locomotor learning (Choi et al., 2009; Darter et al., 2017; Dietz et al., 1995; Morton and Bastian, 2006; Reisman et al., 2007; Roemmich et al., 2014; Roper et al., 2019; Vasudevan et al., 2014) and to design new rehabilitation approaches for persons post-stroke (Reisman et al., 2013, 2010).

Gait adaptation is arguably most useful when learning transfers beyond the practice setting. For example, it is not so valuable that a patient can learn an improved gait pattern during treadmill training; we ultimately want the new walking pattern to transfer overground. It is unclear how the nervous system transfers adaptive learning between contexts, as adapted gait patterns transfer in some situations and not in others. People transfer learning from split-belt walking to overground walking (Reisman et al., 2009; Torres-Oviedo and Bastian, 2010) and between bouts of split-belt walking with different conditions (Leech et al., 2018; Malone et al., 2011) but not between walking and running (Ogawa et al., 2012) or between forward and backward walking (Choi and Bastian, 2007). Features of the training itself – including practice structure (Torres-Oviedo and Bastian, 2012), walking speeds (Vasudevan and Bastian, 2010), and available sensory information (Torres-Oviedo and Bastian, 2010) – also affect transfer. Moreover, movement may not be required at all for learning to occur, as people can learn new reaching movements by watching others adapt their movements (Mattar and Gribble, 2005).

Here, we asked whether people could learn and transfer a new walking pattern without walking. We addressed this question through a series of experiments. In Experiments 1 and 2, we asked whether a new walking pattern could be learned by 1) training each leg independently, or 2) training only one leg. Prior work has shown that stepping with only one leg (i.e., unilateral stepping) drives an aftereffect that persists into normal walking (Huynh et al., 2014; Kahn and Hornby, 2009). This informed our hypothesis that the relative difference in the motion of the legs – not necessarily features of a particular walking pattern, per se (e.g., leg speed) – drives locomotor learning via adaptation. We expected that people could indeed learn a new walking pattern by training the legs unilaterally. In our third experiment, we asked whether active movement was necessary for locomotor learning. Given prior observations in reach adaptation (Mattar and Gribble, 2005; McGregor et al., 2016; Williams and Gribble, 2012), we expected that people could also learn a new walking pattern by watching someone else adapt their gait.

## RESULTS

We performed three experiments to explore different ways that people may be able to learn and transfer a new walking pattern without walking. All experiments took place in a three-dimensional motion capture laboratory where participants walked on a split-belt treadmill (Figure 1A). The experiments followed a similar general protocol (Figure 1B). First, participants walked on the treadmill with both belts moving slowly (0.7 m/s) for two minutes (baseline). Participants then underwent a ten-minute pre-learning period that varied among experiments (Figures 1C-1F). We measured aftereffects of the pre-learning period during a subsequent ten-second catch trial where both belts moved at 0.7 m/s (catch). Participants then walked for ten minutes with the right belt moving at 1.4 m/s and the left moving at 0.7 m/s (learning). This split-belt learning period allowed us to measure transfer of learning from the pre-learning task to a novel walking pattern (i.e., whether the pre-learning task led to faster learning of the split-belt walking pattern). All experiments concluded with the participants walking with both belts at 0.7 m/s for ten minutes to measure aftereffects of learning (post-learning).

**Figure 1.**
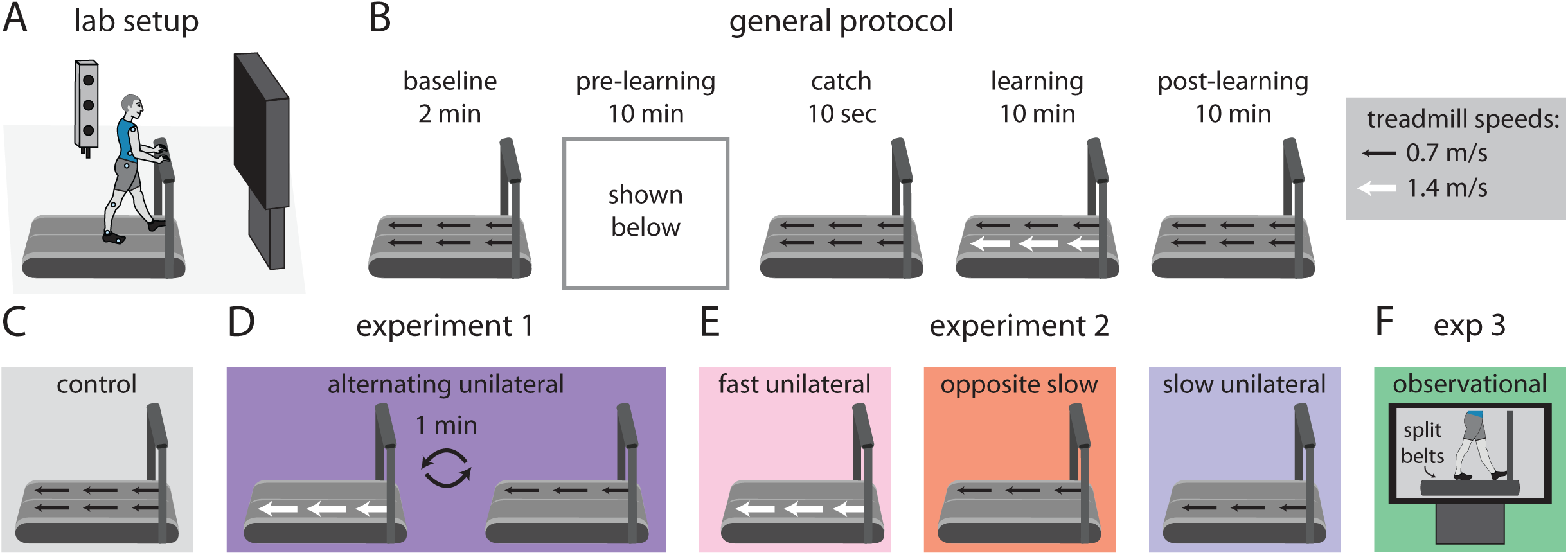
A) Lab setup including split-belt treadmill and three-dimensional motion capture. B) General protocol consisting of 2 minutes of baseline walking, a 10-minute pre-learning condition, 10-second catch trial, 10-minute split-belt learning period, and 10-minute post-learning period. C) Pre-learning in the Control group consisted of walking with both belts moving slowly (0.7 m/s). D) The Alternating Unilateral group in Experiment 1 experienced alternating one-minute bouts of unilateral stepping (right = 1.4 m/s and left = 0 m/s, then right = 0 m/s and left = 0.7 m/s) during pre-learning. E) The Fast Unilateral, Opposite Slow, and Slow Unilateral groups comprised Experiment 2. The pre-learning conditions were: Fast Unilateral (right =1.4 m/s, left = 0 m/s), Opposite Slow (right = 0 m/s, left = 0.7 m/s), and Slow Unilateral (right = 0.7 m/s, left = 0 m/s). F) In Experiment 3, the Observational group watched a 10-minute video of another person adapting their gait to split-belt treadmill walking during pre-learning.

We designed each experiment to answer a specific question. In Experiments 1 and 2, we asked whether training each leg independently or training only one leg, respectively, could drive learning and transfer of a new walking pattern. In Experiment 3, we asked whether people could learn a new walking pattern by observing someone else adapt their gait. We compared the groups from each experiment to a control group using a one-way ANOVA with Dunnett’s post hoc tests (the reference group was the Control group from Experiment 1).

### Experiment 1: can people learn a new walking pattern by training each leg independently?

In Experiment 1, we tested two groups of healthy young adults (n=12 per group). The groups differed only in the tasks performed during the pre-learning period: the Control group walked for ten minutes with both belts moving at 0.7 m/s (Figure 1C) while the Alternating Unilateral group underwent alternating one-minute bouts of unilateral stepping (Figure 1D). During the alternating unilateral stepping, the right treadmill belt moved at 1.4 m/s while the left belt was stationary for one minute, and then the left belt moved at 0.7 m/s while the right belt was stationary for one minute. This alternating pattern repeated five times for a total of ten minutes of training during pre-learning. We chose these speeds because they matched the speeds that each leg would experience during the subsequent split-belt learning period, except here only one leg stepped at a time. We show representative data for the motions of the right and left feet (shown by anterior-posterior ankle marker position) during the Alternating Unilateral pre-learning period in Supplementary Figure 1A.

During the split-belt learning period, the Alternating Unilateral group learned faster than the Control group (Figure 2A), revealing that the movement patterns learned during pre-learning affected the participants’ gait patterns during split-belt walking. We calculated step length asymmetry during the initial perturbation (mean of strides 1-5 during learning), early change (mean of strides 6-30), and plateau (mean of last 30 strides) (Roemmich and Bastian, 2015). The Alternating Unilateral group showed significantly smaller step length asymmetry during early change (p=0.011), indicating significantly faster learning (Figure 2A).

**Figure 2.**
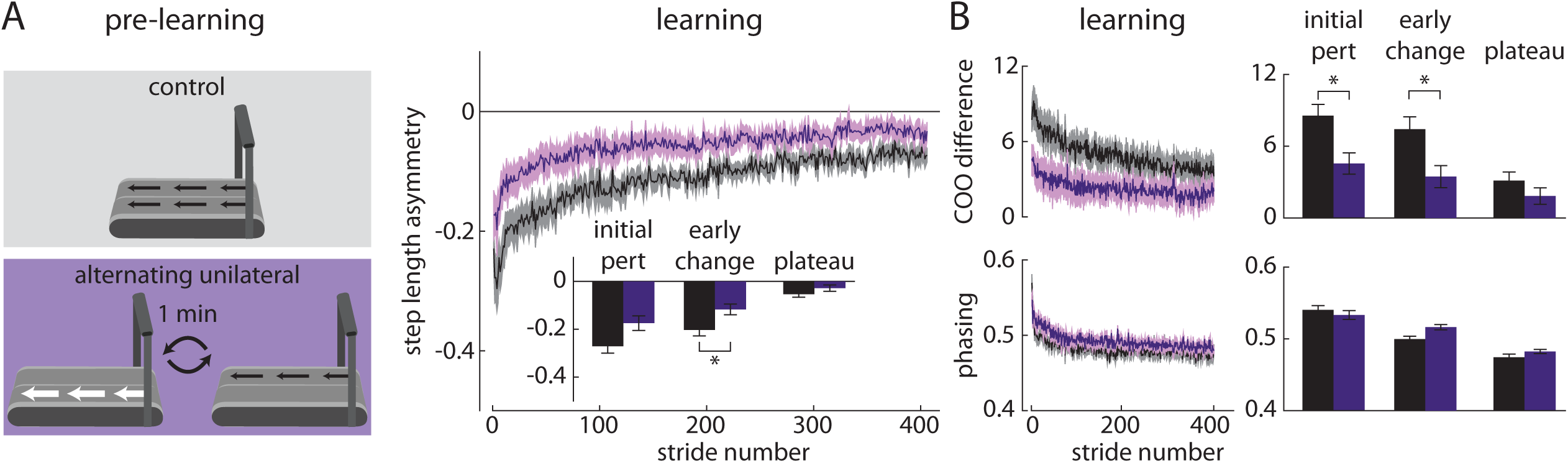
Experiment 1 results. A) The Alternating Unilateral group (purple) showed significantly faster learning (as shown by faster changes in step length asymmetry) during the split-belt learning period when compared to the Control group (black). B) The Alternating Unilateral group also showed significantly faster spatial learning (as shown by faster changes in center of oscillation (COO) difference; top) but similar temporal learning (as shown by phasing; bottom) when compared to the Control group. * indicates p<0.05. In all figures, time series plots are truncated to the length of the data of the participant with the fewest strides.

This faster change in step length asymmetry during learning co-occurred with faster changes in spatial gait parameters (Figure 2B). The Alternating Unilateral group showed significantly smaller center of oscillation (COO) difference during both initial perturbation (p=0.014) and early change (p=0.005) when compared to the Control group (Figure 2B, top). Briefly, COO indicates how each leg oscillates relative to the body. A positive value indicates that the leg oscillates about a flexed axis (i.e., the leg is more often in front of the body); a negative value indicates oscillation around an extended axis (Malone and Bastian, 2010). We determined the spatial symmetry of the gait pattern by finding the COO difference between the legs (0 indicates spatial symmetry). We determined the temporal symmetry of the gait pattern by calculating the phasing between the movements of the legs (Choi and Bastian, 2007). There were no significant differences in phasing between the groups during initial perturbation, early change, or plateau (all p>0.05).

We show data for the baseline, catch, and post-learning periods in Supplementary Figure 2A. We did not observe any significant differences between the groups in step length asymmetry during baseline, catch, or post-learning (all p>0.05). The findings of Experiment 1 showed that people can indeed learn and transfer new walking patterns via adaptation by training each leg independently.

### Experiment 2: can people learn a new walking pattern by training only one leg to step?

We next considered two possible explanations for the results of Experiment 1. First, during pre-learning, the participants may have learned about the speeds that each leg would eventually walk during the split-belt period (“speed match hypothesis”, Figure 3A) and were able to learn and transfer the appropriate movement patterns for each leg. Alternatively, the participants may have learned about the relative motion between the legs – not the speeds of each leg individually – and transferred this learning to split-belt walking (“speed difference hypothesis”, Figure 3B). As shown in Figure 3, these hypotheses make different predictions that we aimed to dissociate in Experiment 2: 1) learning about the leg speeds would accelerate split-belt learning if the legs were first trained to walk at the eventual split-belt speeds (Figure 3A), or 2) learning about the relative motion between the legs would accelerate split-belt learning if the leg that eventually walks faster during split-belt learning also moved faster (i.e., was the moving leg) during unilateral stepping (Figure 3B).

**Figure 3.**
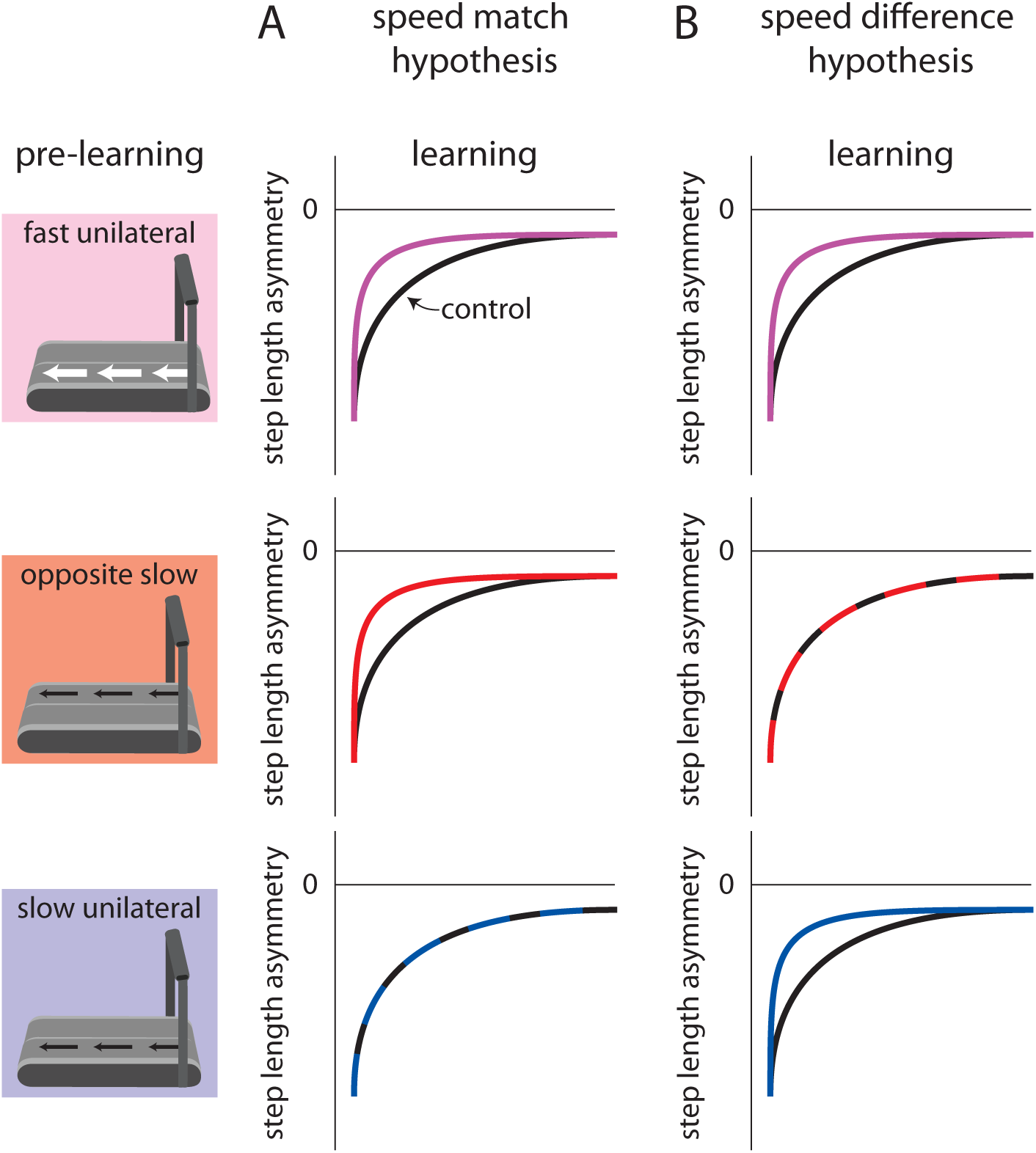
Conceptual figure showing the competing speed match and speed difference hypotheses. These hypotheses make the following dissociable predictions that were tested in Experiment 2: A) the speed match hypothesis predicts faster split-belt learning following Fast Unilateral and Opposite Slow (but not Slow Unilateral) pre-learning conditions where the belt speeds of the moving leg match one of the belt speeds during the subsequent split-belt learning period, and B) the speed difference hypothesis predicts faster split-belt learning following the Fast Unilateral and Slow Unilateral (but not Opposite Slow) pre-learning conditions where the belt that moves during unilateral stepping becomes the faster belt during the subsequent split-belt learning period.

We tested three groups of healthy adults in Experiment 2 (n=12 per group) and compared these groups to the Control group of Experiment 1. First, we considered that both hypotheses predict that a group stepping with the right leg at 1.4 m/s while the left leg was stationary for ten minutes during pre-learning (i.e., the Fast Unilateral group; Figure 1E, left) should replicate our findings from Experiment 1 and show faster learning during the split-belt learning period. This is because 1) the right leg speed during pre-learning matches the right leg speed during learning and, 2) the leg that steps during pre-learning also walks faster during learning (Figure 3A). We indeed observed faster split-belt learning in the Fast Unilateral group. The Fast Unilateral group showed significantly smaller step length asymmetry during initial perturbation (p=0.006) and early change (p<0.001) compared to the Control group (Figure 4A). Similar to the Alternating Unilateral group in Experiment 1, the Fast Unilateral group showed faster changes in spatial (COO difference in Fast Unilateral vs. Control: initial perturbation, p=0.003; early change, p<0.001) but not temporal (phasing: all p>0.05) gait parameters (Figure 4B). We show representative data for the motions of the right and left feet during the Fast Unilateral pre-learning period in Supplementary Figure 1B.

**Figure 4.**
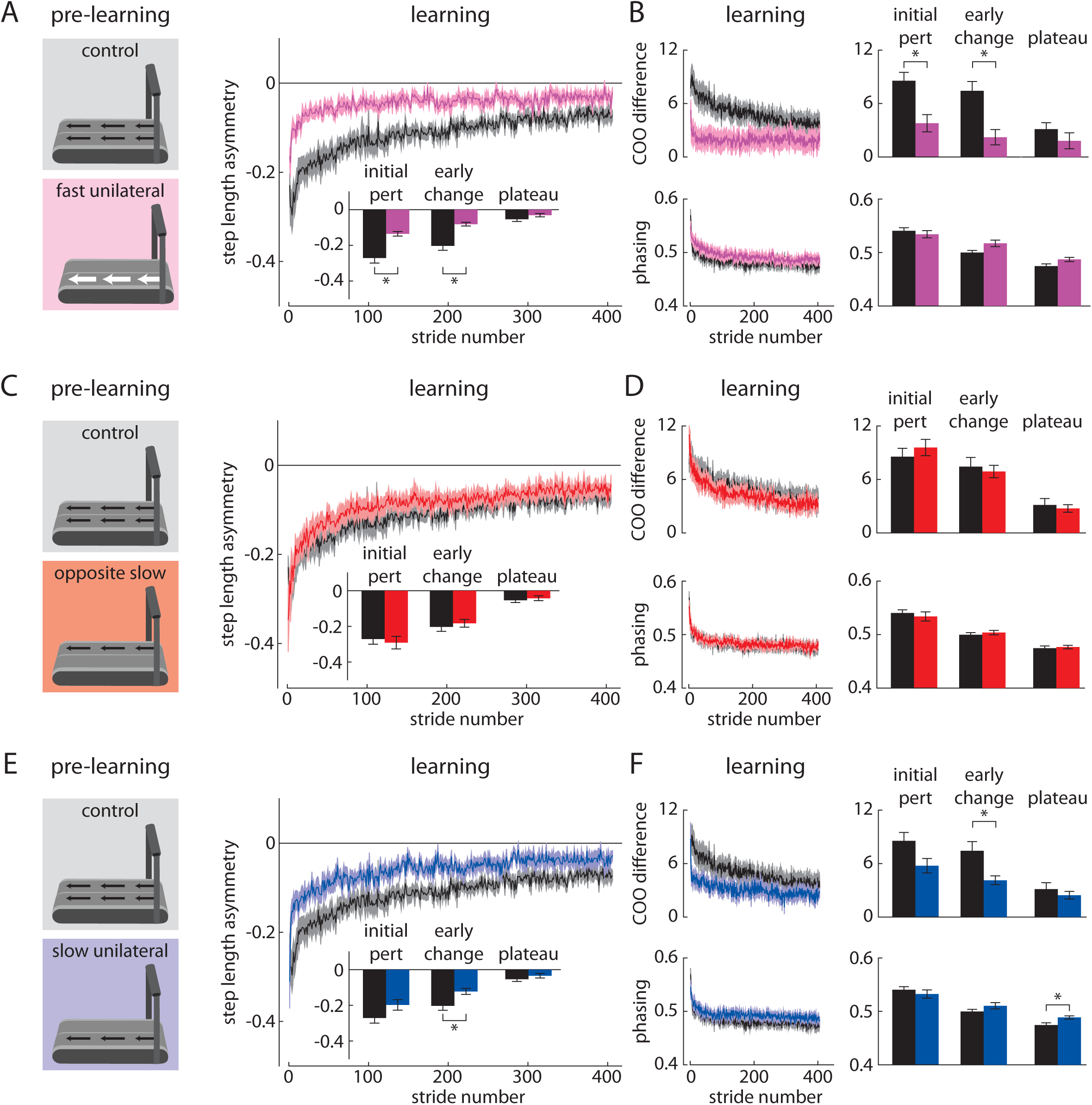
Experiment 2 results. A) The Fast Unilateral group (pink) showed significantly faster learning during the split-belt learning period when compared to the Control group (black). B) The Fast Unilateral group also showed significantly faster spatial learning (top) but similar temporal learning (bottom) when compared to the Control group. C and D) The Opposite Slow group (red) showed similar learning during the split-belt learning period when compared to the Control group. E) The Slow Unilateral group (blue) showed significantly faster learning during the split-belt learning period when compared to the Control group. F) The Slow Unilateral group also showed significantly faster spatial learning (top) but similar temporal learning (except for a small but statistically significant difference during the plateau; bottom) when compared to the Control group. * indicates p<0.05.

We show data for the baseline, catch, and post-learning periods in Supplementary Figure 2B. As expected based on prior findings (Huynh et al., 2014; Kahn and Hornby, 2009), we observed a significant aftereffect in the catch trial following the pre-learning period in the Fast Unilateral group (p<0.001; Supplementary Figure 2B). We did not observe significant differences between the groups in step length asymmetry during baseline or post-learning (all p>0.05).

Next, we aimed to dissociate the competing speed match and speed difference hypotheses. To test the speed match hypothesis, we tested an Opposite Slow group that stepped with the left leg at 0.7 m/s (the same speed that the left leg eventually walked during the split-belt learning period) with the right leg stationary for ten minutes during pre-learning (Figure 1E, middle). In this group, despite the consistent left belt speed across pre-learning and learning, we did not observe faster learning during the learning period (Figure 3A). Instead, as predicted by the speed difference hypothesis, we observed that learning was similar between the Opposite Slow and Control groups (Figure 3B). We did not observe significant differences in step length asymmetry during initial perturbation, early change, and plateau between groups (all p>0.05; Figure 4C). Similarly, we did not observe significant between-group differences in COO difference or phasing (all p>0.05; Figure 4D). We show representative data for the motions of the right and left feet during the Opposite Slow pre-learning period in Supplementary Figure 1C.

We show data for the baseline, catch, and post-learning periods in Supplementary Figure 2C. We observed a significant aftereffect in the catch trial following the pre-learning period in the Opposite Slow group (p=0.004; Supplementary Figure 2C) and a significantly smaller aftereffect during post-learning (initial perturbation, p=0.015) but no other significant differences between the groups in step length asymmetry during baseline or post-learning (all p>0.05).

As a second test of the competing hypotheses, we tested a Slow Unilateral group that stepped with the right leg at 0.7 m/s (the same belt speed difference that the participants eventually experienced during the split-belt learning period but without either of the matched speeds) while the left leg was stationary for ten minutes during pre-learning (Figure 1E, right). As predicted by the speed difference hypothesis (Figure 3B), we observed faster learning during the split-belt learning period. The Slow Unilateral group showed significantly smaller step length asymmetry during early change (p=0.018) compared to the Control group (Figure 4E). Similar to the Alternating Unilateral and Fast Unilateral groups, the Slow Unilateral group showed faster changes in spatial (COO difference in Slow Unilateral vs. Control: early change, p=0.024) but not temporal (phasing: initial perturbation and early change, p>0.05) gait parameters (Figure 4F), though there was a significant difference in the phasing plateau (p=0.024). We show representative data for the motions of the right and left feet during the Slow Unilateral pre-learning period in Supplementary Figure 1D.

We show data for the baseline, catch, and post-learning periods in Supplementary Figure 2D. We observed a significant aftereffect in the catch trial following the pre-learning period in the Slow Unilateral group (p=0.046; Supplementary Figure 2D) that was similar in magnitude and opposite in direction to that of the Opposite Slow group (Supplementary Figure 2C), as would be expected. We did not observe significant differences between the groups in step length asymmetry during baseline or post-learning (all p>0.05).

### Does the amount learned during unilateral stepping predict faster learning during split-belt walking?

Given that step length asymmetry during the catch trial reveals how much was learned during the pre-learning period, we considered that step length asymmetry during the catch trial should predict step length asymmetry during the learning period if learning had truly transferred from pre-learning to learning. In other words, if a participant learned more during pre-learning, and learning transfers from the pre-learning period to the learning period, the participant should show faster learning during the learning period. We performed two stepwise linear regressions – one with step length asymmetry during initial perturbation of the split-belt learning period as the dependent variable and one with step length asymmetry during early change as the dependent variable, both with group assignment variables and step length asymmetry during the catch trial as independent variables – to test the relationship between pre-learning and learning across all groups in Experiment 2. We did not include the Alternating Unilateral group from Experiment 1 because we considered that the catch trial aftereffect may have disappeared due to a savings effect from the switching between right and left unilateral stepping conditions, and thus this analysis was limited to groups that only stepped with one leg during pre-learning. We found that step length asymmetry during the catch trial was indeed a significant predictor of step length asymmetry during initial perturbation (standardized β=0.62, p<0.001; Figure 5, top) and early change (standardized β=0.64, p<0.001; Figure 5, bottom) while the group assignment variables were not (all p>0.05).

**Figure 5.**
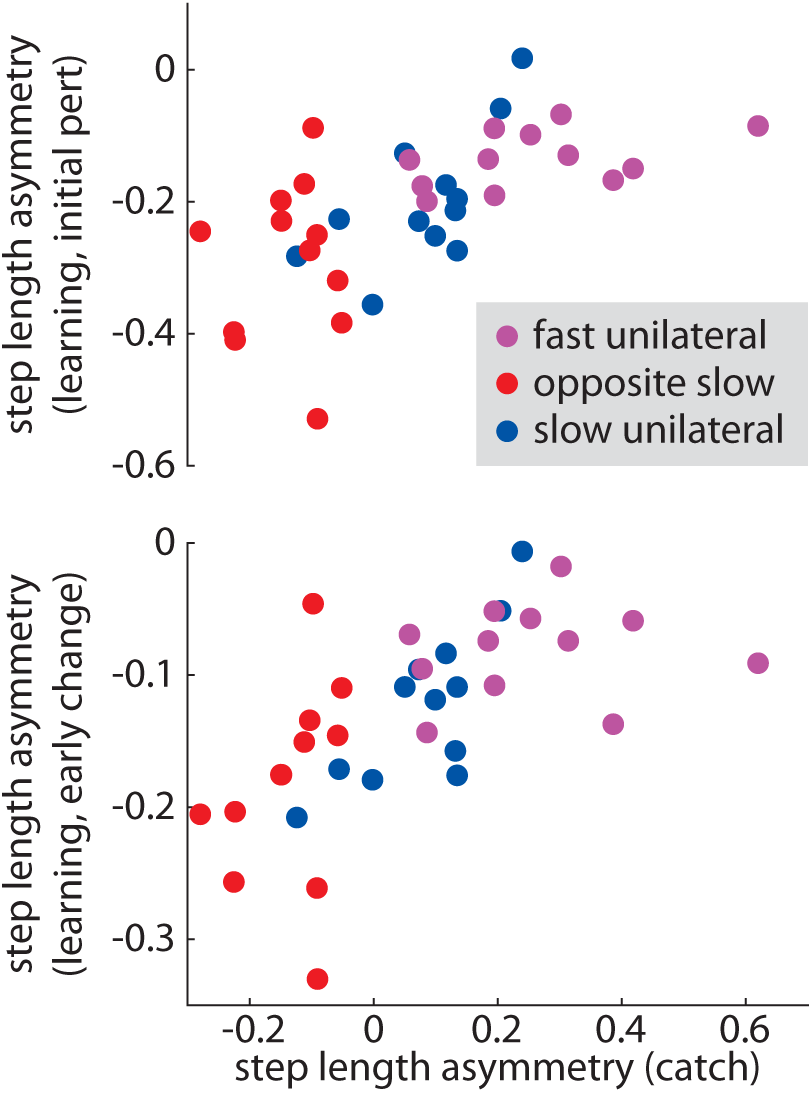
Across the groups in Experiment 2, step length asymmetry measured during the catch trial (i.e., the pre-learning aftereffect) was a significant predictor of the step length asymmetry during initial perturbation (top) and early change (bottom) during split-belt learning (i.e., learning rate). Colors are consistent with prior figures. Regression β and p values are included in the Results section.

The results of Experiment 2 revealed that people could learn and transfer a new walking pattern by training only one leg to step. These findings support the hypothesis that the nervous system learned about the relative motion between the legs during the pre-learning unilateral stepping. Transfer then occurred into the split-belt learning period, and this accelerated learning was accompanied by accelerated changes in spatial gait parameters. Our findings did not support the hypothesis that the nervous system learns and transfers information about the individual leg speeds.

### Experiment 3: can people learn a new walking pattern by observing someone else adapt their gait?

Experiments 1 and 2 showed that, by engaging in specific patterns of unilateral step training, people could learn and transfer a new walking pattern without bilateral gait training. In Experiment 3, we asked whether active movement was necessary for locomotor learning at all. We considered that people may be able to learn a new walking pattern by simply watching someone else adapt their gait (Mattar and Gribble, 2005). In Experiment 3, we tested an Observational group (n=12) that watched a ten-minute video of a naïve participant learning to walk on a split-belt treadmill (sagittal view, waist-down) during pre-learning (Figure 1F). Contrary to our hypothesis, we did not observe significantly faster learning in the Observational group (as compared to the Control group) during the split-belt learning period. We did not observe significant differences between the groups in step length asymmetry during initial perturbation, early change, or plateau (all p>0.05; Figure 6A). COO difference and phasing were also similar between groups (all p>0.05; Figure 6B). We also did not observe a significant aftereffect in the catch trial following the pre-learning period (p>0.05; Supplementary Figure 2E). We did observe a significantly smaller aftereffect during post-learning (initial perturbation, p=0.036) but did not observe significant differences between the groups in step length asymmetry during baseline or post-learning (all p>0.05). In Experiment 3, we conclude that people did not learn the new walking pattern significantly faster after watching someone else adapt their gait.

**Figure 6.**
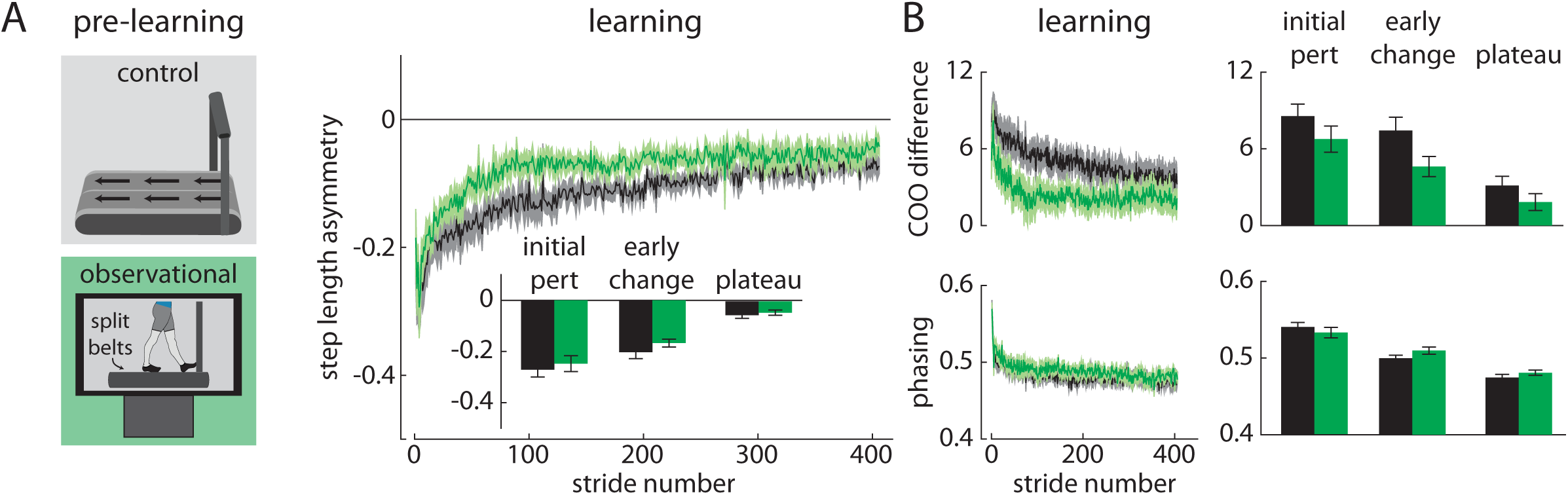
Experiment 3 results. A and B) The Observational group (green) showed similar learning during the split-belt learning period when compared to the Control group (black).

### Do changes in spatial gait parameters predict faster learning across all groups?

Several of our analyses in Experiments 1-3 showed that changes in step length asymmetry during learning co-occurred with changes in spatial gait parameters (e.g., COO difference). To explore the relationships between changes in step length asymmetry and changes in spatial (and temporal) gait parameters during learning more directly, we performed two additional stepwise linear regressions that included all data from Experiments 1-3. In the first regression, the dependent variable was step length asymmetry during initial perturbation of the split-belt learning period. The independent variables were group assignment variables, COO difference during initial perturbation, and phasing during initial perturbation. Only COO difference was a significant predictor of step length asymmetry during initial perturbation (standardized β=-0.85, p<0.001; Figure 7, top left); phasing (Figure 7, top right) and group assignment variables were not. In the second regression, the dependent variable was step length asymmetry during early change of the split-belt learning period. The independent variables were group assignment variables, COO difference during early change, and phasing during early change. Only COO difference (standardized β=-0.82, p<0.001; Figure 7, bottom left) and assignment to the Fast Unilateral group (standardized β=-0.15, p=0.032) were significant predictors of step length asymmetry during early change; phasing (Figure 7, bottom right) and other group assignment variables were not. These findings revealed that changes in spatial gait parameters strongly predicted changes in step length asymmetry during learning (i.e., learning rate).

**Figure 7.**
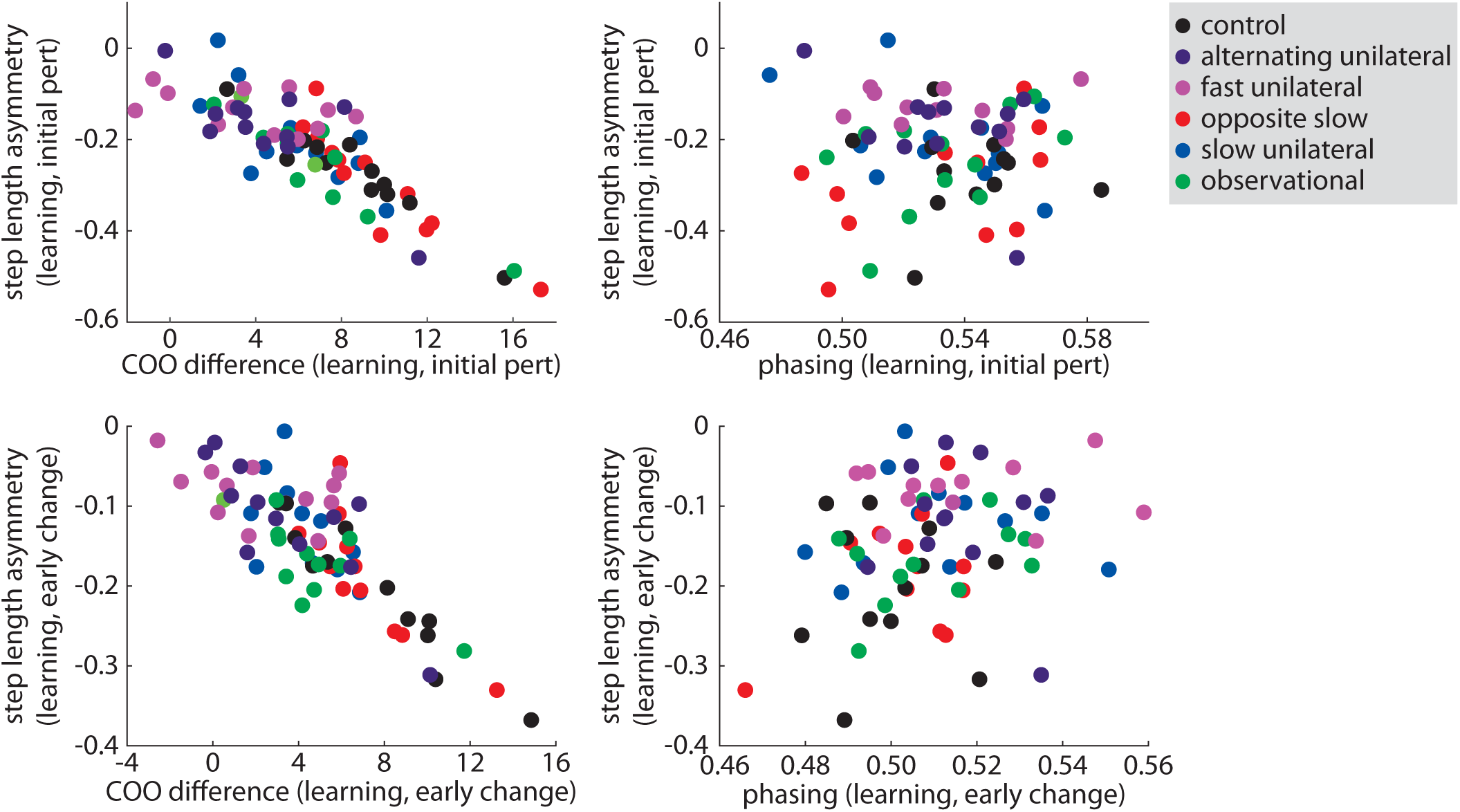
Across all groups in Experiments 1-3, COO difference (i.e., spatial learning) was a significant predictor of step length asymmetry during initial perturbation (top left) and early change (bottom left) during split-belt learning. Phasing (i.e., temporal learning) was not a significant predictor of step length asymmetry during initial perturbation (top right) or early change (bottom right) during split-belt learning. Colors are consistent with prior figures. Regression β and p values are included in the Results section.

## DISCUSSION

The results of this study demonstrated that people could learn and transfer a new walking pattern without walking. We found that training only one leg to step at a time could drive transfer of learning that resulted in accelerated acquisition of a novel walking pattern. Our findings also revealed that the nervous system learns and transfers information about the relative motion of the legs – not the appropriate movement pattern for a specific speed – during adaptive learning, regardless of whether both legs are in motion. In summary, we have shown that people can learn new walking patterns without bilateral gait training, as unilateral adaptation-based approaches can result in learning that transfers to novel gait patterns.

A key finding of this study is that unilateral step training only accelerated split-belt learning under certain conditions. Split-belt learning occurred faster following unilateral stepping when the stepping leg eventually walked on the faster belt during the split-belt learning condition. Several prior studies offer potential mechanisms that could explain why this occurs. Adaptation is thought to be a cerebellar-mediated motor learning process driven by sensory prediction error (Martin et al., 1996; Morton and Bastian, 2006; Tseng et al., 2007). While it is not clear what constitutes the prediction error during split-belt treadmill walking, the repeated exposures to similar patterns of belt speed differences (between unilateral stepping and split-belt walking conditions) likely caused similar prediction errors. Many studies have shown that learning occurs faster when people experiences similar errors repeatedly (Herzfeld et al., 2014; Kojima et al., 2004; Leech et al., 2018; Malone et al., 2011; Medina et al., 2001; Roemmich and Bastian, 2015), a phenomenon termed ‘savings’ (or ‘retention’, if the learning periods are not separated by a washout period). Savings also occurs when the first perturbation is larger than the second (Leech et al., 2018), a finding that aligns with our observations in the Fast Unilateral group. It is likely that the faster split-belt learning observed in the Alternating Unilateral, Fast Unilateral, and Slow Unilateral groups was due to the nervous system perceiving similar errors during the pre-learning and learning periods and facilitating a retention-like response. This also explains why the belt speed difference – not the individual belt speeds – was the key variable driving transfer.

We showed that transfer of spatial learning drives the faster transfer of step length asymmetry seen with unilateral stepping in this study. COO difference is a spatial measure of gait asymmetry that reflects that one leg takes steps that are placed in front of the steps by the other leg. The directionality of this change (i.e., which leg steps further forward than the other) is determined by the difference between the belt speeds in split-belt treadmill walking or unilateral stepping: the leg on the faster belt learns to step farther ahead than the slower leg. In this way, the important role of spatial learning in mediating transfer of unilateral stepping agrees with the idea that it is the belt speed difference, not individual belt speeds, that determines transfer. Our observation that faster changes in step length asymmetry were mediated by faster spatial learning is consistent with several prior studies of savings in split-belt treadmill walking (Malone et al., 2011; Musselman et al., 2016; Roemmich and Bastian, 2015) and hybrid stepping where one leg marches in place while the other walks normally (Long et al., 2015). The relative difference in placements of the feet at heel-strike may be a key variable that the nervous system uses in order to learn and facilitate transfer.

Why does spatial gait asymmetry (COO difference), but not temporal gait asymmetry (phasing) explain the faster transfer of step length asymmetry with unilateral stepping? Several studies have suggested that spatial and temporal gait asymmetry are controlled by different processes in the nervous system (Choi et al., 2009; Darmohray et al., 2019; Gonzalez-Rubio et al., 2019; Malone et al., 2012). Temporal gait asymmetry has previously been shown to adapt faster than spatial asymmetry during split-belt treadmill walking (Malone et al., 2012), an observation that our results support. Our results show that temporal gait asymmetry during learning was unaffected by unilateral stepping or observational learning during pre-learning. This invariance may be explained by a recent study that suggests a hierarchical organization that prioritizes corrections to temporal asymmetry over spatial asymmetry (Gonzalez-Rubio et al., 2019). Thus, temporal gait asymmetry may be controlled via processes distinct from the ‘error-based’ mechanism that governs spatial learning.

The finding that unilateral stepping can accelerate learning of novel gait patterns may benefit rehabilitation of individuals who cannot tolerate prolonged walking bouts. For these individuals, unilateral stepping may be a training modality that is tolerable and confers benefits that not only transfer to overground walking (Kahn and Hornby, 2009) but facilitate learning of new walking patterns. Furthermore, unilateral stepping is accessible, as it requires only a conventional treadmill (a patient can stand with one leg to the side of the treadmill and step with the contralateral leg). Prior work established that unilateral stepping induces changes in step length asymmetry during subsequent overground walking in healthy adults (Huynh et al., 2014) and improves step length asymmetry in persons with chronic hemiparesis (Kahn and Hornby, 2009). Our results suggest that unilateral stepping can accelerate learning of a novel gait pattern through a similar error-based adaptation mechanism that is normally associated with locomotor learning on a split-belt treadmill (Reisman et al., 2010). Adaptive learning appears to be a promising mechanism for facilitating transfer of learning from unilateral to bilateral stepping.

Unlike what has been shown in experiments of reaching adaptation (Mattar and Gribble, 2005; McGregor et al., 2018, 2016; Williams and Gribble, 2012), we did not observe a significant effect of observational learning on locomotor adaptation. Though the comparisons between the Observational and Control group did not reach statistical significance, we noted modestly faster learning in the Observational group that occurred later in the learning curve. The metrics in our study focused on changes over the first 30 strides (i.e., where the largest changes in the curve tend to occur), and the Observational group appears to separate modestly from the Control group beyond this time epoch. It is possible that observational learning may transfer slightly to locomotor adaptation, though this may occur through more explicit learning instead of the error-based learning that we believe transfers following the unilateral stepping conditions.

In addition to studying the transfer from the pre-learning to learning conditions, we also assessed aftereffects of unilateral stepping or observation (pre-learning) to tied-belt walking (catch trial) and aftereffects of split-belt treadmill (learning) to tied-belt walking (post-learning). Unilateral stepping produced aftereffects in the catch trial that were likely mediated through a speed difference mechanism. For example, the aftereffects observed in the catch trials following the Fast Unilateral, Opposite Slow and Slow Unilateral conditions showed magnitudes and directions that changed in accordance with the speed difference of the preceding unilateral stepping condition. One potentially puzzling finding is that, in contrast to the aftereffects observed in the Fast Unilateral, Opposite Slow and Slow Unilateral conditions, the Alternating Unilateral group showed no aftereffect during the catch trial. We suggest that this may be driven by savings achieved from switching repeatedly between the two unilateral stepping patterns in the Alternating Unilateral condition. As mentioned above, switching between two conditions repeatedly eventually leads to rapid changes in walking patterns (Malone et al., 2011), and this can be explained by a computational model of motor adaptation that accelerates learning by increasing the sensitivity to errors upon repeated exposures to the same errors (Herzfeld et al., 2014). Future work that improves our understanding of the prediction errors experienced during unilateral stepping and split-belt walking (particularly with regard to the belt speed difference that drives the learning) may provide additional insight.

There are some features of our data that we found more difficult to explain. It is unclear why the aftereffect disappears in the catch trial following Alternating Unilateral stepping but reappears in the same group following split-belt walking. If we think that unilateral stepping and split-belt walking learn from similar errors, one expects a smaller aftereffect following split-belt walking in this group (as was observed in the Opposite Slow group). We also lack explanations for why the aftereffect following split-belt walking was smaller in the Observational group than the Control group given that there were no significant differences in any other conditions between the groups (though there appears to be a qualitative difference between these groups during learning, it did not reach statistical significance) and why there is a small but statistically significant difference in temporal learning plateau between the Slow Unilateral and Control groups.

Lastly, we think it is important to clarify that the pre-learning conditions studied here are different than the pre-gait activities incorporated in some rehabilitation approaches. Pre-gait activities have been used for patients deemed not yet ready for gait training and promote practice of specific gait features (e.g., weight shifting, limb advancement) outside of a locomotor setting. This approach relies on repetitive training over time and is minimally effective at improving walking in neurologic populations (Eng and Tang, 2007). Conversely, we have shown that training the legs independently of one another through an adaptation-based approach transferred to a novel gait pattern in healthy young participants. While this approach may have promising implications for teaching patients new gait patterns, we emphasize that our results do not suggest that this training approach should be used as a precursor to task-specific gait training to promote the recovery of walking function. Our findings also do not suggest that all unilateral training approaches (regardless of learning mechanism) should be expected to result in effective locomotor learning.

## CONCLUSION

Locomotor adaptation can be leveraged to drive people to learn new walking patterns without walking. We have shown that training the legs individually or only training one leg to step on a treadmill can facilitate learning of a novel gait pattern. The accelerated learning observed in some groups during split-belt walking was mediated by transfer of spatial learning from preceding unilateral stepping conditions. This was influenced by the difference between the leg speeds experienced in unilateral stepping; in contrast, the legs did not learn to move at a particular speed in isolation of one another, nor did we observe evidence of observational learning. Adaptive learning can be leveraged to drive transfer of movement patterns learned via unilateral stepping to novel bilateral gait patterns.

## MATERIALS AND METHODS

### Participants

Seventy-two healthy young adults participated (mean±SEM age: 20.4±0.2 years, sex: 41 F/31 M). All participants were naïve to split-belt treadmill walking and unilateral stepping. Participants were free of neurological, musculoskeletal, or cardiovascular conditions and participated in only one experiment. All participants provided written informed consent in accordance with the Johns Hopkins Medicine Institutional Review Board prior to participation. They were compensated monetarily for their participation and watched a television show or movie of their choice during the experiment (except during the pre-learning period in the Observational group).

### Data Collection

We used an Optotrak Certus motion capture system (100 Hz; Norther Digital, Waterloo, ON) to collect kinematic data as participants walked (or stepped unilaterally) on a split-belt treadmill (Woodway USA, Waukesha, WI). We recorded kinematic data from active markers placed bilaterally over the fifth metatarsal head, lateral malleolus, lateral femoral epicondyle, greater trochanter, iliac crest, and acromion. We controlled the belt speeds using custom MATLAB software (MathWorks, Natick, MA). A thin partition approximately 12 inches tall ran lengthwise between the belts to prohibit stepping on both belts simultaneously. This did not otherwise interfere with walking. We notified participants when the treadmill was about to start or stop moving but did not tell them how the treadmill would move except for the unilateral stepping trials, where we told participants that only one belt would move and they were to keep the opposite foot on the ground at all times. Participants held onto the handrails as the treadmill accelerated but released them immediately thereafter. We stopped the treadmill briefly between conditions (e.g., baseline, pre-learning, catch, learning, post-learning), and participants remained on the treadmill across all conditions. Participants wore comfortable shoes, form-fitting clothing, and a safety harness that did not provide body weight support.

### Experimental protocols

As shown in Figure 1B and described in the Results, all experiments followed a similar general protocol. Briefly, experiments began by collecting a ten-second trial of quiet standing. Participants then walked with both belts moving at the same speed (0.7 m/s, baseline) followed by a ten-minute pre-learning task that differed among groups:

- Control: both belts at 0.7 m/s
- Alternating Unilateral: alternating one-minute periods where the right belt moved at 1.4 m/s and left was stationary or the right was stationary and the left moved at 0.7 m/s
- Fast Unilateral: right at 1.4 m/s, left 0 m/s
- Opposite Slow: right 0 m/s, left at 0.7 m/s
- Slow Unilateral: right at 0.7 m/s, left 0 m/s
- Observational: both belts stationary as participants stood and watched a ten-minute video of a waist-down, sagittal view of a naïve participant adapting to split-belt treadmill walking

We then recorded a ten-second catch trial where both belts moved at 0.7 m/s to measure pre-learning aftereffects (catch). After the catch trial, participants walked under split-belt conditions where the right and left belts moved at 1.4 m/s and 0.7 m/s, respectively, for ten minutes (learning). Finally, participants again walked with both belts at 0.7 m/s to measure learning aftereffects (post-learning).

### Data analysis

Our primary outcome measure was step length asymmetry. We calculated step lengths for each leg as the anterior-posterior distance between the lateral malleolus (i.e., ankle) markers at heel-strike and then calculated step length asymmetry for each stride as:

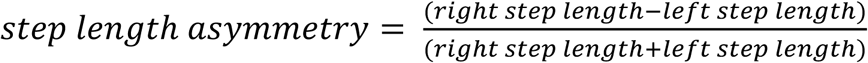

Because both legs are in constant motion during treadmill walking, step length is influenced by both spatial (where the foot is placed) and temporal (when the foot is placed relative to the contralateral foot) control. Therefore, step length asymmetry is a measure of spatiotemporal gait symmetry. We also calculated metrics of spatial (COO difference) and temporal (phasing) symmetry. These metrics have been well-described elsewhere (Choi and Bastian, 2007; Malone and Bastian, 2010) and are described briefly in the Results. COO difference is a measure of the difference between the axes around which each leg oscillates during walking, and phasing is a measure of the temporal coordination (e.g., how in-phase or out-of-phase) of the oscillations of each leg.

We calculated the mean step length asymmetry, COO difference, and phasing during several time periods. In the baseline and catch periods, we calculated the means of each metric across the entire period for each participant. In the learning and post-learning periods, we calculated the means during the initial perturbation (strides 1-5), early change (strides 6-30), and plateau (last 30 strides) epochs for each participant. We did not calculate these metrics during pre-learning because only one leg stepped at a time in a majority of the groups (i.e., the metrics were not possible to calculate during unilateral stepping).

### Statistical analysis

We used one-way ANOVA with Dunnett’s post hoc tests to assess differences in our primary (step length asymmetry) and secondary (COO difference, phasing) outcome measures across groups. All groups in Experiments 1-3 were included in the analysis, and the Control group from Experiment 1 was treated as the reference group for the post hoc tests. We compared our outcome measures across groups in the baseline, catch, learning (initial perturbation, early change, and plateau), and post-learning (initial perturbation, early change, and plateau) periods.

We also performed two pairs of stepwise linear regressions as described in the Results. In the first pair of regressions, the dependent variables were step length asymmetry during initial perturbation of the split-belt learning period and one with step length asymmetry during early change. Both included group assignment variables and step length asymmetry during the catch trial as independent variables. We performed these regressions to test the relationship between pre-learning and learning across all unilateral stepping groups in Experiment 2. In the second pair of regressions, the dependent variables were step length asymmetry during the initial perturbation and early change epochs of the split-belt learning period. The independent variables were 1) group assignment variables, COO difference during initial perturbation, and phasing during initial perturbation, and 2) group assignment variables, COO difference during early change, and phasing during early change. We performed these regressions to test the relationship between changes in step length asymmetry and changes in spatial (COO difference) and temporal (phasing) symmetry during the split-belt learning period across all groups.

We performed all statistical analyses using SPSS (IBM, Armonk, NY) and used an alpha level of 0.05.

## ACKNOWLEDGMENTS

This study was funded by NIH grant R21 AG059184.

## FIGURE LEGENDS

**Supplementary Figure 1.**
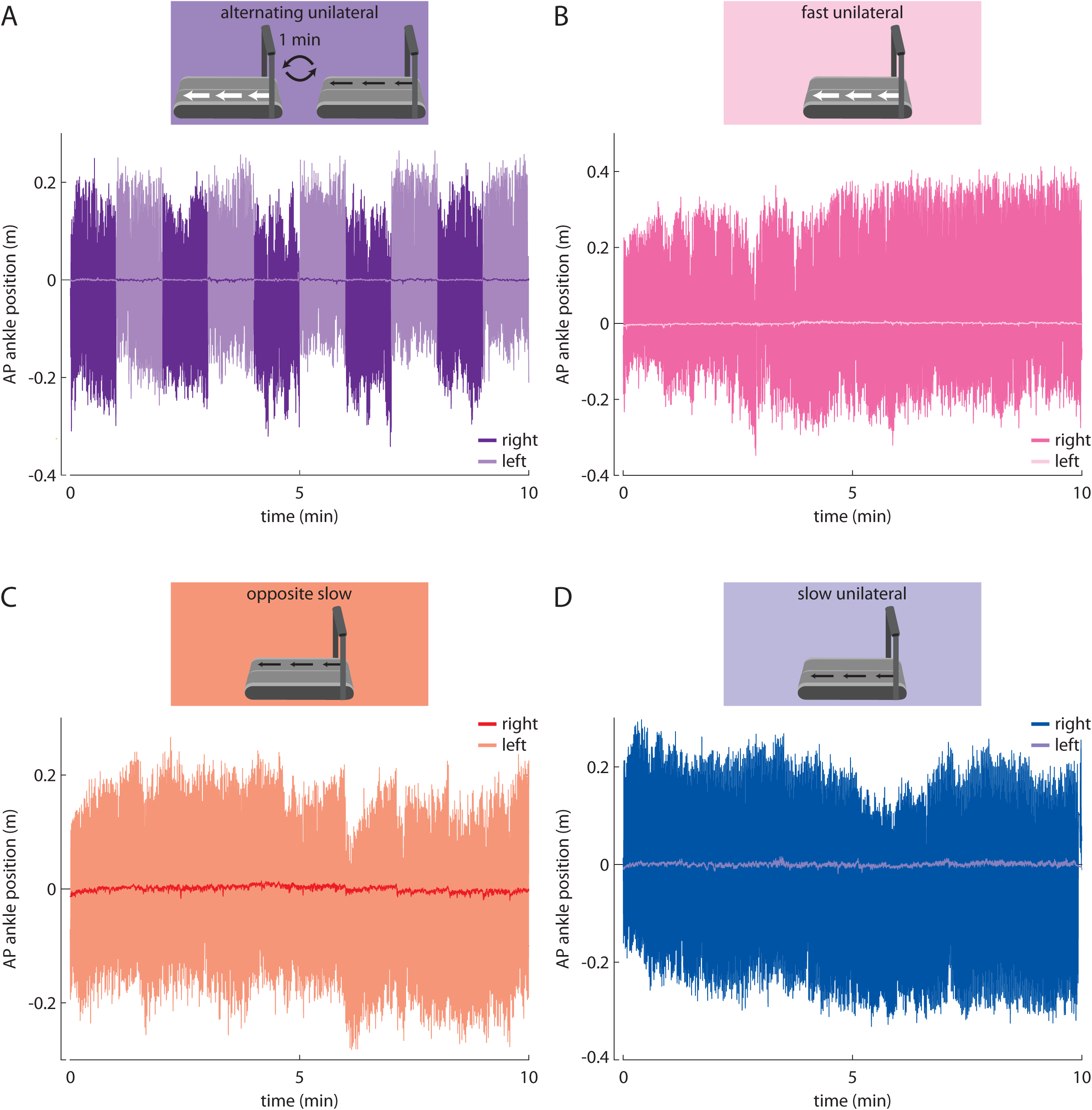
Representative individual examples of foot movement (i.e., anterior-posterior ankle marker position) during the pre-learning conditions in the A) Alternating Unilateral, B) Fast Unilateral, C) Opposite Slow, and D) Slow Unilateral groups. Darker lines indicate right leg, lighter lines indicate left leg.

**Supplementary Figure 2.**
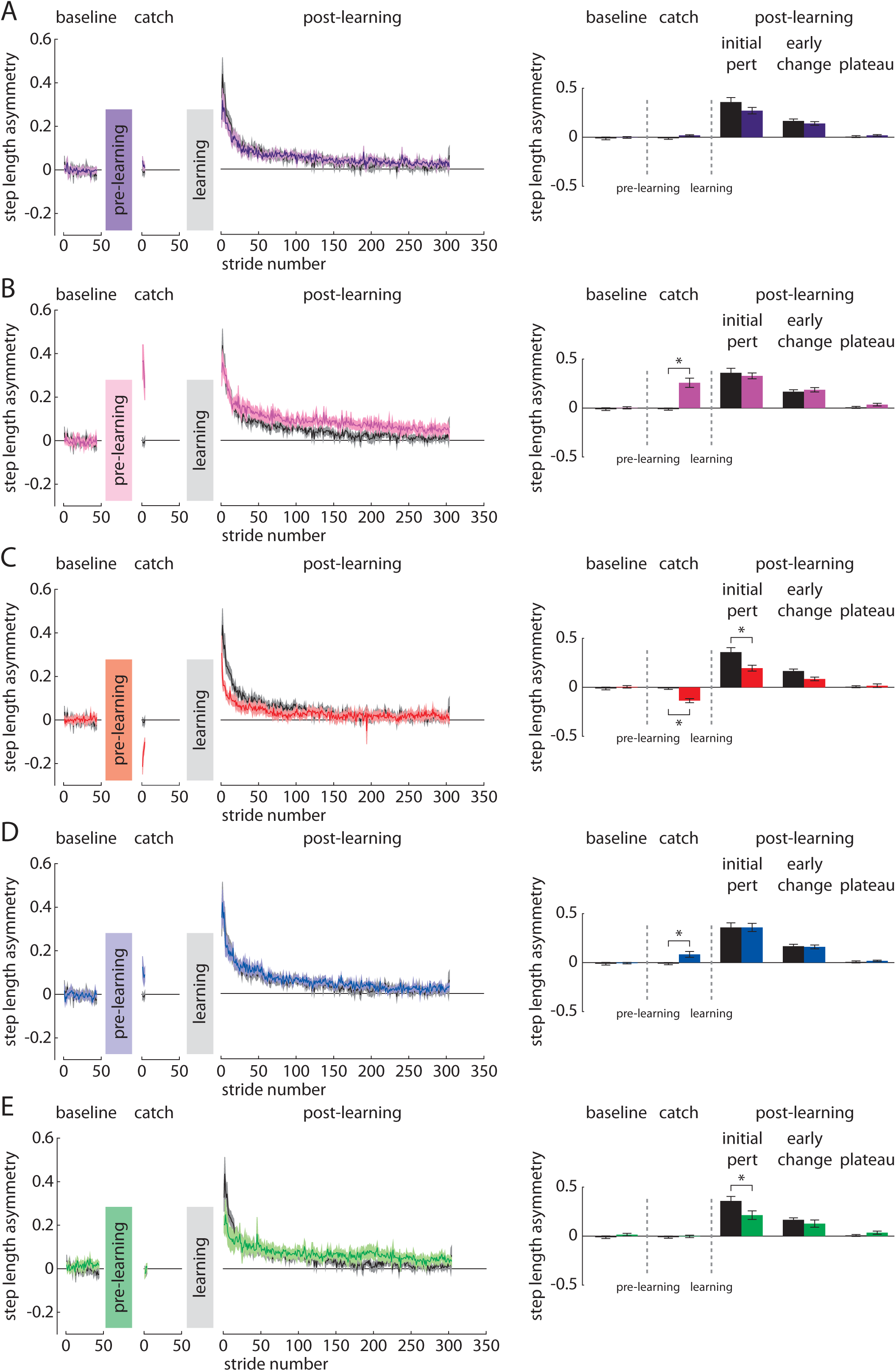
Step length asymmetry during the baseline, catch, and post-learning periods in the A) Alternating Unilateral (purple), B) Fast Unilateral (pink), C) Opposite Slow (red), D) Slow Unilateral (blue), and E) Observational (green) groups compared to Control (black). * indicates p<0.05.

